# A single loss of photosynthesis in diatoms

**DOI:** 10.1101/298810

**Authors:** Anastasiia Onyshchenko, Elizabeth C. Ruck, Teofil Nakov, Andrew J. Alverson

## Abstract

Loss of photosynthesis is a common and often repeated trajectory in nearly all major groups of photosynthetic eukaryotes. One small subset of ‘apochloritic’ diatoms in the genus *Nitzschia* have lost their ability to photosynthesize and require extracellular carbon for growth. Similar to other secondarily nonphotosynthetic taxa, apochloritic diatoms maintain colorless plastids with highly reduced plastid genomes. Although the narrow taxonomic breadth of apochloritic diatoms suggests a single loss of photosynthesis in the common ancestor of these species, previous phylogenetic analyses suggested that photosynthesis was lost multiple times. We sequenced additional phylogenetic markers from the nuclear and mitochondrial genomes for a larger set of taxa and found that the best trees for datasets representing all three genetic compartments provided low to moderate support for monophyly of apochloritic *Nitzschia*, consistent with a single loss of photosynthesis in diatoms. We sequenced the plastid genome of one apochloritic species and found that it was highly similar in all respects to the plastid genome of another apochloritic *Nitzschia* species, indicating that streamlining of the plastid genome had completed prior to the split of these two species. Finally, it is increasingly clear that some locales host relatively large numbers apochloritic *Nitzschia* species that span the phylogenetic diversity of the group, indicating that these species co-exist because of resource abundance or resource partitioning in ecologically favorable habitats. A better understanding of the phylogeny and ecology of this group, together with emerging genomic resources, will help identify the factors that have driven and maintained the loss of photosynthesis in this group, a rare event in diatoms.

## Introduction

Photosynthetic eukaryotes (Archaeplastida) trace back to a single common ancestor, in which a eukaryotic host paired with a cyanobacterial endosymbiont that would eventually become the plastid, a fully integrated cellular organelle that is the site of photosynthesis (Archibald 2009, Keeling 2010). Although most archaeplastids remain photoautotrophic, loss of photosynthesis has occurred—often repeatedly—in nearly all major archaeplastid lineages (Hadariová et al. 2017). As many as five different lineages within the green algal orders Chlamydomonadales and Chlorellales have traded off photosynthesis for trophic strategies that include heterotrophy and obligate parasitism (Rumpf et al. 1996, Tartar and Boucias 2004, Yan et al. 2015). Partial or complete loss of photosynthesis has occurred in at least 11 different lineages of flowering plants, representing hundreds of species (Barkman et al. 2007); parasitic angiosperms obtain extracellular carbon from the vascular tissues of host plants for all (holoparasites) or part (hemiparasites) of their life history (Těšitel 2016). Perhaps most strikingly, photosynthesis has been lost dozens or more times across the florideophycean red algae (e.g., Goff et al. 1996, Kurihara et al. 2010), which go on to parasitize a closely related photosynthetic species—a phenomenon known as adelphoparasitism (Blouin and Lane 2012). Photosynthesis has been lost in a broad range of taxa with secondary plastids as well, including euglenoids (Marin 2004), apicocomplexans (McFadden et al. 1996, Waller and McFadden 2005), ciliates (Reyes-Prieto et al. 2008), dinoflagellates (Saldarriaga et al. 2001), cryptophytes (Donaher et al. 2010, Martin-Cereceda et al. 2010), and stramenopiles (Tyler et al. 2006). Most nonphotosynthetic algae evolved from mixotrophic ancestors (Figueroa-Martinez et al. 2015), likely because they already had the means to secure extracellular carbon and because the energetic costs of mixotrophy are thought to be high (Raven 1997).

Diatoms are a lineage of stramenopile algae responsible for roughly 20% of global primary production (Field et al. 1998). They are ancestrally photosynthetic, and although the overwhelming majority of the estimated 100,000 or so diatom species remain photosynthetic, many species are mixotrophic, which allows them to use external sources of carbon for growth in fluctuating light conditions (Hellebust and Lewin 1977, Tuchman et al. 2006). A much smaller set of 20 or so mostly undescribed, colorless, free-living species in the genus *Nitzschia*, and one species in the closely related and morphologically similar genus Hantzschia, have abandoned photosynthesis altogether and rely exclusively on extracellular carbon for growth (Lewin and Lewin 1967). These “apochloritic” diatoms—the only known nonphotosynthetic diatom species—are often found in association with mangroves, and decaying seaweeds and sea grasses (Pringsheim 1956, Blackburn et al. 2009a, Kamikawa et al. 2015b).

The small number of species and narrow taxonomic range of apochloritic diatoms leads to the prediction, based on parsimony, that nonphotosynthetic diatoms are monophyletic, tracing back to a single loss of photosynthesis in their common ancestor. Despite the small number of species, however, they encompass a relatively broad range of morphological diversity (Blackburn et al. 2009b, Kamikawa et al. 2015b) and use a variety of carbon sources (Lewin and Lewin 1967, Hellebust and Lewin 1977), raising the possibility that obligate heterotrophy evolved multiple times—a hypothesis supported by a phylogenetic analysis of nuclear 28S d1–d2 sequences that separated apochloritic taxa into three separate clades (Kamikawa et al. 2015b). Monophyly of apochloritic species could not be rejected, however (Kamikawa et al. 2015b). These two competing hypotheses (one vs. multiple origins) have important implications for our understanding of the underlying phylogenetic, ecological, and genomic contexts of this radical trophic shift, which has occurred far less frequently in diatoms than it has in other groups.

Like most other lineages that have lost photosynthesis, apochloritic diatoms maintain highly reduced plastids and plastid genomes (Kamikawa et al. 2015a). Their plastids lack chlorophyll and thylakoids (Kamikawa et al. 2015b), and their plastid genomes have lost most photosynthesis-related genes, including all photosystem genes (Kamikawa et al. 2015a). The nearly complete set of ATP synthase genes in the plastid genome might function in ATP hydrolysis, creating a proton gradient that fuels protein import into the plastid (Kamikawa et al. 2015a). Carbon metabolism is highly compartmentalized in diatoms (Smith et al. 2012), and the large number of nuclear-encoded proteins targeted to the plastid point to a highly metabolically active and, as a result, indispensable (Kamikawa et al. 2017) organelle. Although comparative genomics is greatly improving our understanding of carbon metabolism in both photosynthetic (Smith et al. 2012) and nonphotosynthetic diatoms (Kamikawa et al. 2015a, 2017), the power of comparative genomics can only be fully leveraged within the framework of an accurate, densely sampled phylogenetic hypothesis.

We collected, isolated, and cultured several apochloritic *Nitzschia* species and sequenced common phylogenetic markers to test whether photosynthesis was lost one or multiple times. A combined dataset of nuclear, mitochondrial, and plastid genes support monophyly of apochloritic *Nitzschia* species, consistent with a single loss of photosynthesis in diatoms. Species were split between two major subclades that co-occur in habitats with apparently favorable, albeit poorly defined, conditions. We also sequenced and characterized the plastid genome of *Nitzschia* sp. and found it to be highly similar to that of another species in the same subclade, indicating rapid genomic streamlining following loss of photosynthesis in their common ancestor. The results presented here help frame a number of questions about the evolution, ecology, and genomic consequences of this rare, radical trophic shift in diatoms.

## Materials and Methods

### Collection and culturing of apochloritic *Nitzschia*

All cultures originated from a single composite sample, collected on 10 November 2011 from Whiskey Creek, which is located in Dr. Von D. Mizell-Eula Johnson State Park (formerly John U. Lloyd State Park), Dania Beach, Florida, USA (26.081330 lat, –80.110783 long). This site was previously shown to host a diverse assemblage of apochloritic *Nitzschia* species (Blackburn et al. 2009a). Our sample consisted of near-surface plankton collected with a 10 *µm* mesh net, submerged sand (1m and 2m depth), and nearshore wet (but unsubmerged) sand. We selected for nonphotosynthetic species by storing the sample in the dark at room temperature (21°C) for several days before isolating colorless diatom cells with a Pasteur pipette. Clonal cultures were grown in the dark at 21°C on agar plates made with L1+NPM medium (Guillard 1960, Guillard and Hargraves 1993) and 1% Penicillin–Streptomycin–Neomycin solution (Sigma-Aldrich P4083) to retard bacterial growth.

### DNA extraction, PCR, and DNA sequencing

Cells were rinsed with L1 medium and removed from agar plates by pipetting, briefly centrifuged, then broken with MiniBeadbeater-24 (BioSpec Products). We then extracted DNA with a Qiagen DNeasy Plant Mini Kit. Nuclear SSU and partial LSU rDNA genes were PCR-amplified and sequenced using published PCR conditions and primer sequences (Alverson et al. 2007). PCR and sequencing primers for two mitochondrial markers, cytochrome b (*cob*) and NADH dehydrogenase subunit 1 (*nad1*), are listed in Table S1. PCRs for mitochondrial genes used: 1.0–5.0 *µL* of DNA, 6.5 *µL* of Failsafe Buffer E (Epicentre Technologies), 0.5 *µL* of each primer (20 *µM* stocks), 0.5 units Taq polymerase, and ddH2O to a final volume of 25 *µL*. In a few cases, we used a nested PCR strategy to amplify the *nad1* gene. PCR conditions for *cob* and *nad1* genes were as follows: 95 °C for 5 minutes, 36 cycles of (95 °C for 60 *s*, 45 °C for 60 *s*, 72 °C for 60 *s*), and a final extension at 72 °C for 5 minutes. PCR products were sequenced on an ABI 3100 capillary sequencer. Raw sequences were assembled and edited with Geneious ver. 7.1.4 (Biomatters Ltd.) and deposited in GenBank (Table 1).

**Table 1.** Taxa and sources of DNA, and GenBank accessions for sequences analyzed in this study. Apochloritic taxa are in bold face.

| Genus | Species | Culture.ID | cob | nad1 | lsu | 16s |
| --- | --- | --- | --- | --- | --- | --- |
| Nitzschia | sp. | ccmp2144 | MH017326 | MH017300 | MH017352 |  |
| Nitzschia | dippelii | AJA014-6 | MH017327 | MH017301 | MH017353 |  |
| Hantzschia | spectabilis | UTEX FD269 | MH017328 | MH017302 | MH017354 |  |
| Hantzschia | elongata | UTEX FD421 | MH017329 | MH017303 | MH017355 |  |
| Hantzschia | amphioxys | AJA013-11 | MH017330 | MH017304 | MH017356 |  |
| Fragilariopsis | cylindrus | ccmp1022 | JGI16035 | JGI16036 |  |  |
| Pseudo-nitzschia | sp. | ccmp1447 | MH017331 | MH017305 | MH017357 |  |
| Nitzschia | frustulum | ccmp1532 | MH017332 | MH017306 | MH017358 |  |
| Nitzschia | cf. ovalis | ccmp1118 | MH017333 | MH017307 | MH017359 |  |
| Cylindrotheca | closterium | ccmp1855 | MG271845 | MG271845 |  |  |
| Psammodictyon | constrictum | ccmp576 | MH017334 | MH017308 | MH017360 |  |
| Nitzschia | laevis | ccmp1092 | MH017335 | MH017309 | MH017361 |  |
| Nitzschia | sp. | OHI12.10 | MH017336 | MH017310 | MH017362 |  |
| Nitzschia | cf. frequens | ccmp1500 | MH017337 | MH017311 | MH017363 |  |
| Nitzschia | pusilla | ccmp2526 | MH017338 | MH017312 | MH017364 |  |
| Nitzschia | cf. pusilla | ccmp581 | MH017339 | MH017313 | MH017365 |  |
| Nitzschia | sp. | ccmp2533 | MH017340 | MH017314 | MH017366 |  |
| Nitzschia | draveiliensis | AJA010-17 | MH017341 | MH017315 | MH017367 |  |
| Nitzschia | palea var. debilis | AJA013-2 | MH017342 | MH017316 | MH017368 |  |
| Nitzschia | gracilis | AJA010-53 | MH017343 | MH017317 | MH017369 |  |
| Nitzschia | capitellata | AJA012-22 | MH017344 | MH017318 | MH017370 |  |
| **Nitzschia** | **alba** | **ccmp2426** | MH017345 | MH017319 | MH017371 |  |
| **Nitzschia** | **leucosigma** | **ccmp2197** | MH017346 | MH017320 | MH017372 | MH017379 |
| **Nitzschia** | **sp.** | **ccmp579** | MH017347 | MH017321 | MH017373 |  |
| **Nitzschia** | **sp.** | **Nitz2** | MH017348 | MH017322 | MH017377 |  |
| **Nitzschia** | **sp.** | **Nitz4** | MH017349 | MH017323 | MH017375 | MH017378 |
| **Nitzschia** | **sp.** | **Nitz7** | MH017350 | MH017324 | MH017376 |  |
| **Nitzschia** | **sp.** | **Nitz8** | MH017351 | MH017325 | MH017374 | MH017380 |

### Phylogenetic analyses

In addition to the data generated here, we compiled previously published data for the plastid 16S and nuclear 28S rDNA genes for Bacillariales to better enable comparisons with previous studies. We downloaded all Bacillariales sequences from Genbank, checked percent identity and coverage of each sequence to a local BLAST database comprised of apochloritic *Nitzschia* sequences, and reverse-complemented sequences if necessary. We kept Bacillariales sequences that were at least 20% identical and covered at least 40% of at least one target in the database (BLAST options: identity cutoff=20, hsp coverage cutoff=40). We kept only the longest sequence for cases in which multiple downloaded sequences had the same NCBI Taxid identifier. The total number of sequences meeting these criteria were 46 for the 16S and 116 for the 28S rDNA genes.

We used SSU-ALIGN ver. 0.1 (Nawrocki et al. 2009) to align rDNA sequences (plastid 16S and the d1–d3 region of the nuclear 28S rDNA), using SSU-ALIGN’s built-in covariance models of secondary structure for bacteria for the 16S alignment and a heterokont-specific covariance model for the 28S alignment (Nakov et al. 2014). We used SSU-MASK to remove poorly aligned regions and used these more conservative alignments for downstream analyses. The protein-coding *cob* and *nad1* genes from the mitochondrial genome were aligned by hand with Mesquite ver. 3.31 (Maddison, W. P. and Maddison, D. R. 2008) after color-coding nucleotide triplets by their conceptual amino acid translations. We built phylogenetic trees from each individual gene alignment and three concatenated alignments: (1) a concatenation of the two mitochondrial genes into a *cob*+*nad1* (heretofore mito dataset), (2) a concatenation of the two mitochondrial genes and the masked 28S alignment of newly generated 28S sequences (heretofore mito-lsu dataset), and (3) a concatenation of the two mitochondrial genes and the masked alignment of all 28S sequences (newly generated and downloaded from GenBank, heretofore mito-ncbi-lsu dataset). The alignments have been archived in a ZENODO online repository (https://doi.org/10.5281/zenodo.1211571).

We used RAxML ver. 8.2.4 (Stamatakis 2014) to infer phylogenetic trees from each of the three concatenated alignments. For each dataset, we performed 10 maximum likelihood searches to find the best-scoring tree and a rapid bootstrap analysis of 250 pseudoreplicates per search (Stamatakis et al. 2008), for a total of 2500 bootstrap samples per alignment. We used the general time-reversible model (GTR) of nucleotide substitution, and we used a Γ distribution to accommodate rate variation across sites within each alignment (GTR+GAMMA, the default model in RAxML).

### Plastid genome sequencing and analysis

We sequenced the plastid genome of *Nitzschia* sp. (Nitz4) using the Illumina HiSeq2000 platform, with a 300-bp library and 90-bp paired-end reads. We removed adapter sequences and trimmed raw reads with Trimmomatic ver. 0.32 and settings ‘ILLUMINACLIP=<TruSeq adapters.fasta>:2:30:10, TRAILING=5, SLIDINGWINDOW=6:18, HEADCROP=9, MINLEN=50’ (Bolger et al. 2014). We assembled trimmed reads using Ray ver. 2.3.1 with default settings and k-mer length of 45 (Boisvert et al. 2012). We assessed the assembly quality with QUAST ver. 2.3 (Gurevich et al. 2013), confirmed high genome-wide read coverage by mapping the trimmed reads to the assembly with Bowtie ver. 0.12.8 (Langmead et al. 2009), and evaluated these results with SAMtools. We used DOGMA (Wyman et al. 2004) and ARAGORN (Laslett and Canback 2004) to annotate the genome. The annotated genome has been archived in GenBank under accession MG273660. We used Easyfig (Sullivan et al. 2011) to perform a BLASTN-based synteny comparison of the plastid genomes of two apochloritic *Nitzschia* taxa, strains Nitz4 (this study) and NIES-3581 (Kamikawa et al. 2015a).

## Results

### Phylogeny of apochloritic *Nitzschia*

We isolated and cultured four strains (Nitz2, Nitz4, Nitz7, and Nitz8) of apochloritic *Nitzschia* from Whiskey Creek (Florida, USA), including both linear and undulate forms (Fig. 1). Although cells grew well in culture, we never observed sexual reproduction and, unlike some other apochloritic *Nitzschia* strains that have been maintained in culture for decades, all of our strains experienced substantial size reduction and eventually died off.

**Figure 1.**
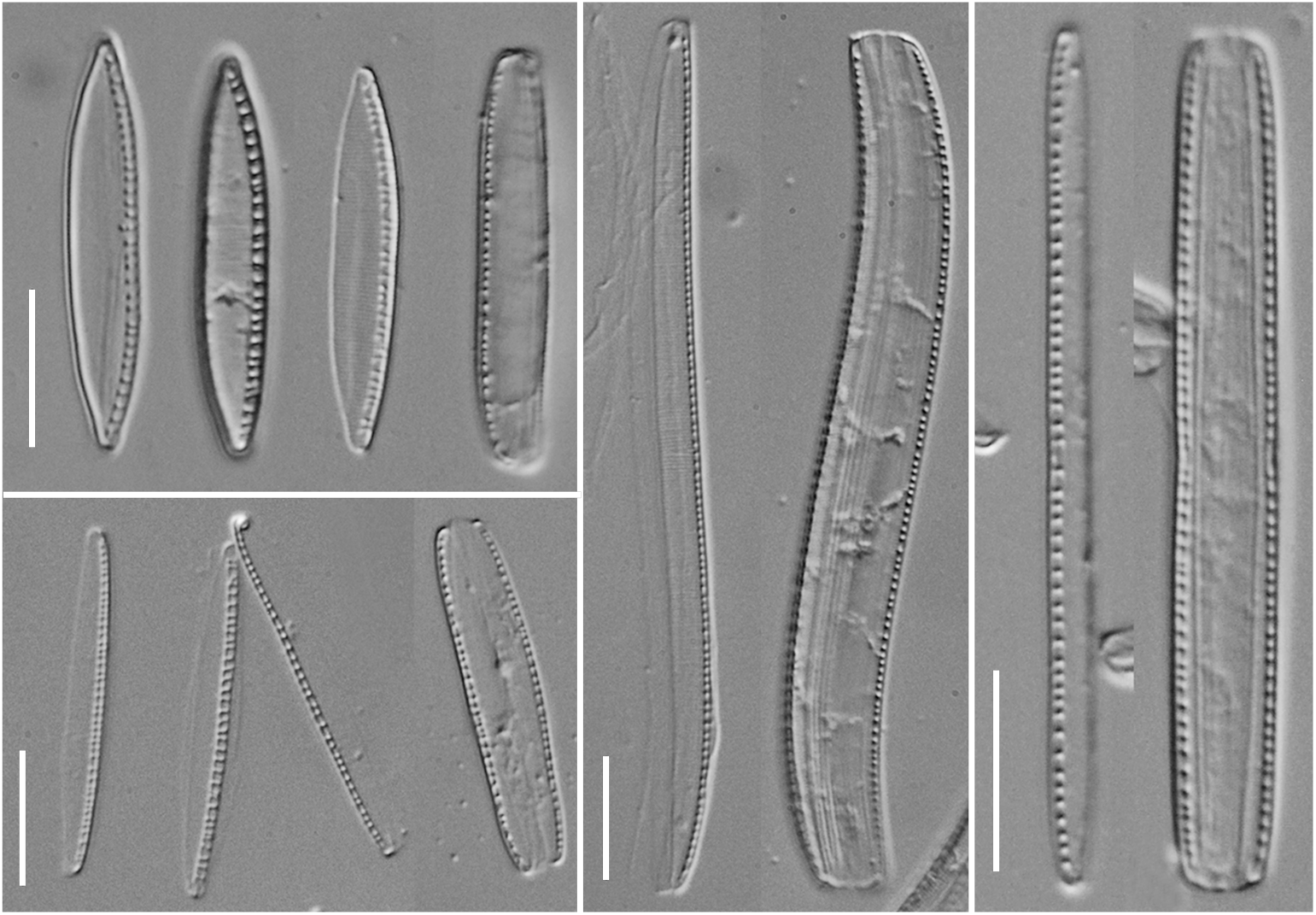
Light micrographs of newly sequenced nonphotosynthetic *Nitzschia* species. Clock-wise from top: *Nitzschia* sp. nitz2, *Nitzschia* sp. nitz7, *Nitzschia* sp. nitz8, *Nitzschia* sp. nitz4.

We sequenced the nuclear 28S rDNA and mitochondrial *cob* and *nad1* genes for 26 Bacillariales taxa, including seven apochloritic strains (Table 1). We combined these data with GenBank sequences to reconstruct phylogenetic trees for individual and concatenated alignments. The plastid 16S rDNA gene supported monophyly of apochloritic *Nitzschia* species (Fig. 2A, Bootstrap proportion (BS)=100), though the exceptionally long branch lengths–driven by decreased GC content–raises the possibility that this grouping simply reflects shared nucleotide composition (Steel et al. 1993, Galtier and Gouy 1995, Kamikawa et al. 2015b). One way to reduce the influence of GC bias in tree inference is to transform the alignment into purine/pyrimidine (R/Y) coding. By treating all A and G nucleotides as R and all C and T nucleotides as Y, the GC bias present in some sequences is masked, the base frequencies are normalized, and the majority of phylogenetic signal originates from transversions (i.e., R ↔ Y). The phylogeny resulting from the R/Y-transformed alignment should better reflect the history of the lineage rather than the nucleotide composition bias. Performing this transformation for the plastid 16S alignment, we again found strong support for monophyly of the apochloritic *Nitzschia* (bootstrap support = 95%; Fig 2B). This result suggests that monophyly of apochloritic *Nitzschia* species reconstructed with the plastid-encoded 16S gene might not necessarily be an artifact of the decreased GC content in the plastid genomes of nonphotosynthetic taxa.

**Figure 2.**
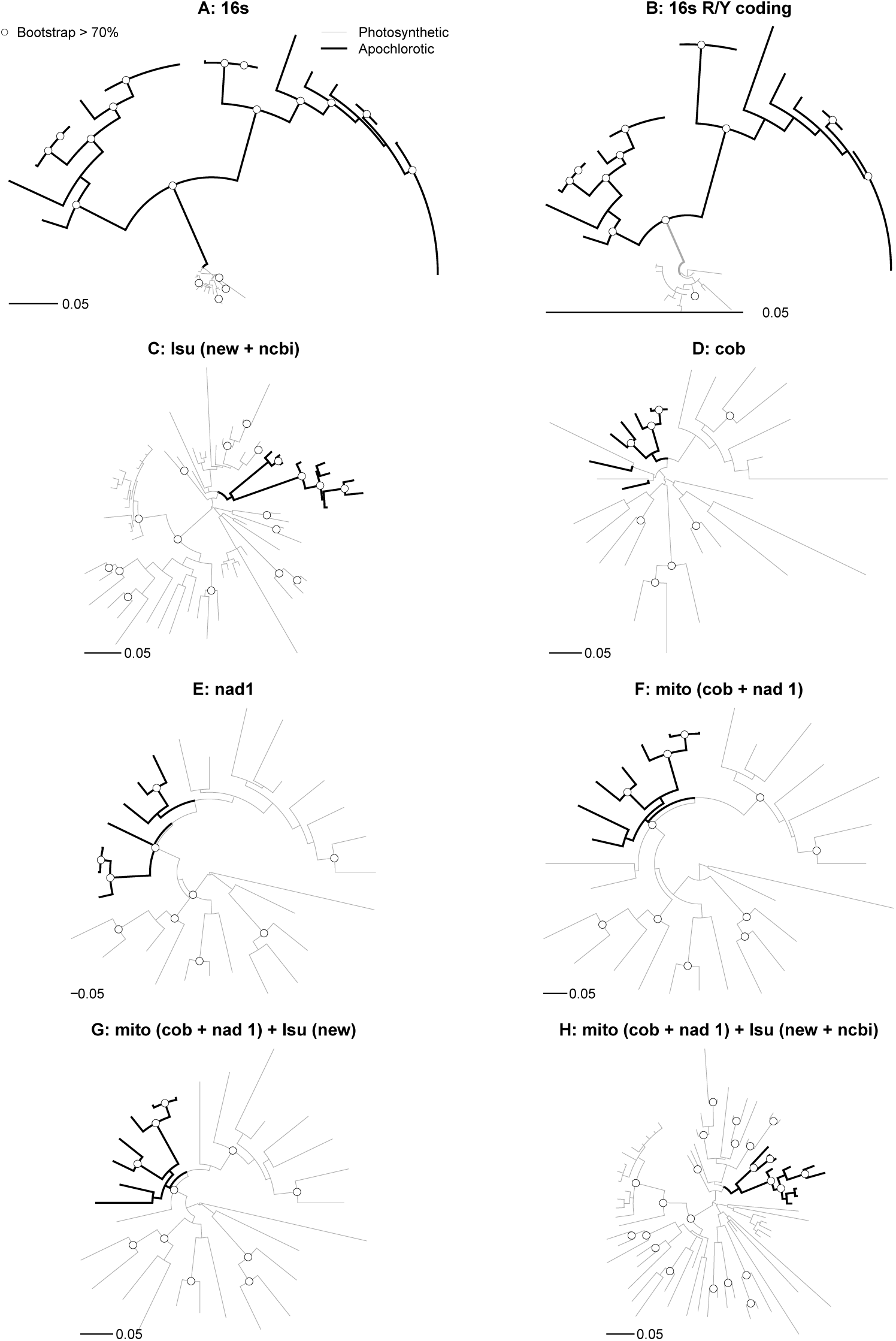
Phylogenetic trees inferred from plastid 16S (A), plastid 16S transformed into purine/pyrimidine (R/Y) coding (B), nuclear 28S d1–d2 genes for all newly sequenced taxa and data from GenBank [”lsu (new + ncbi)”, C], mitochondrial *cob* (D), mitochondrial *nad1* (E), concatenated *cob* and *nad1* genes [”mito (*cob* + *nad1*)”, F], concatenated *cob*, *nad1*, and nuclear 28S d1–d2 genes for all newly sequenced taxa [”mito (*cob* + *nad1*) + lsu (new)”, G], and a concatenated alignment of *cob*, *nad1*, and nuclear 28S d1–d2 genes for taxa from this study and GenBank [”mito (*cob* + *nad1*) + lsu (new + ncbi)”, H]. For the phylogenies in panels C and H, we removed branches shorter than 0.00001 units for clarity. The full phylogenies are available in Supplementary Figure 1. Thicker black branches correspond to apochlorotic taxa. White points identify nodes with bootstrap support *>*70%.

A previous analysis of nuclear 28S d1–d2 rDNA sequences weakly supported non-monophyly of apochloritic taxa, though monophyly could not be rejected (Kamikawa et al. 2015b). Expanding the analysis to include the large number of Bacillariales 28S d1–d2 sequences from GenBank and newly sequenced taxa from this study (Table 1) returned a monophyletic grouping of apochloritic *Nitzschia* (Fig. 2C). We also sequenced two mitochondrial genes, *cob* and *nad1*, and although neither of them individually supported monophyly of apochloritic taxa (Fig. 2D, E), analysis of a concatenated mitochondrial *cob*+*nad1* alignment did return a clade of apochloritic *Nitzschia* (Fig. 2F). Concatenated alignments of nuclear 28S d1–d2 and the two mitochondrial genes supported monophyly of apochloritic *Nitzschia* as well, both for an analysis restricted to just newly sequenced taxa (Fig. 2G) and one that included both newly sequenced and GenBank taxa (Fig. 2H). In all cases, branch support for monophyly of apochloritic *Nitzschia* with nuclear and mitochondrial markers, analyzed individually or in combination, was less than 70% (Fig. 2). The fully labeled trees are available in Fig. S1.

### Plastid genome sequencing and analysis

The plastid genome of *Nitzschia* sp. Nitz4 maps as a circular chromosome and has the conserved quadripartite architecture typical of most plastid genomes, with two inverted repeat (IR) regions and small and large single copy regions (SSC and LSC, respectively). At 67,764 bp in length, the genome is roughly half the size of plastid genomes from photosynthetic diatoms (Ruck et al. 2014, Yu et al. 2018). The genome contains minimal intergenic DNA, with 12 genes that overlap by anywhere from 1–61 bp in length and another two genes that are immediately adjacent to one another. (Table S2). The genome consists mainly of genes with conserved housekeeping functions, including 32 tRNA and six rRNA genes. All three rRNA and six of the tRNA genes are present twice in the genome because of their location in the IR region. More than half (45) of the 73 protein-coding genes encode ribosomal proteins or subunits of RNA polymerase, with ATP synthase genes and ORFs constituting most of the remaining genes. Some of the ORFs include ones (ycf41, ycf89 and ycf90) that are highly conserved among diatoms (Ruck et al. 2014, Yu et al. 2018). The genome also contains Sec-independent protein translocator TatC and and SecA subunit of Sec-mediated transport system, iron-sulfur clusters (SufB and SufC), molecular chaperones dnaK, and a protease (ClpC). The genome is highly AT-rich (77.6% A+T).

Similar to the plastid genome of another apochloritic diatom, *Nitzschia* sp. NIES-3581 (aka IriIs04) (Kamikawa et al. 2015a), the genome of *Nitzschia* sp. Nitz4 is missing all genes encoding subunits of photosystem I and II, proteins of cytochrome b6f complex, carbon fixation system, porphyrin and thiamine metabolism. As well Nitz4 lacks several conserved ORFs and membrane translocators subunits, dnaB helicase, chaperonine groEL and ftsH cell division protein. The genomes of Nitz4 and NIES-3581 are highly similar in size, gene content, gene order, nucleotide composition, and sequence (Fig. 3). The loss of rps20 in Nitz4, which appears to have been replaced by a unique ORF (orf122), is among the few minor differences between the two genomes (Fig. 3). Two gene fusions, orf96-atpB and orf122-rpoB, present in Nitz4 are separated in NIES-3581. Likewise, the overlapping dnaK-tRNA-Arg genes in NIES-3581 are adjacent but non-overlapping in Nitz4.

**Figure 3.**
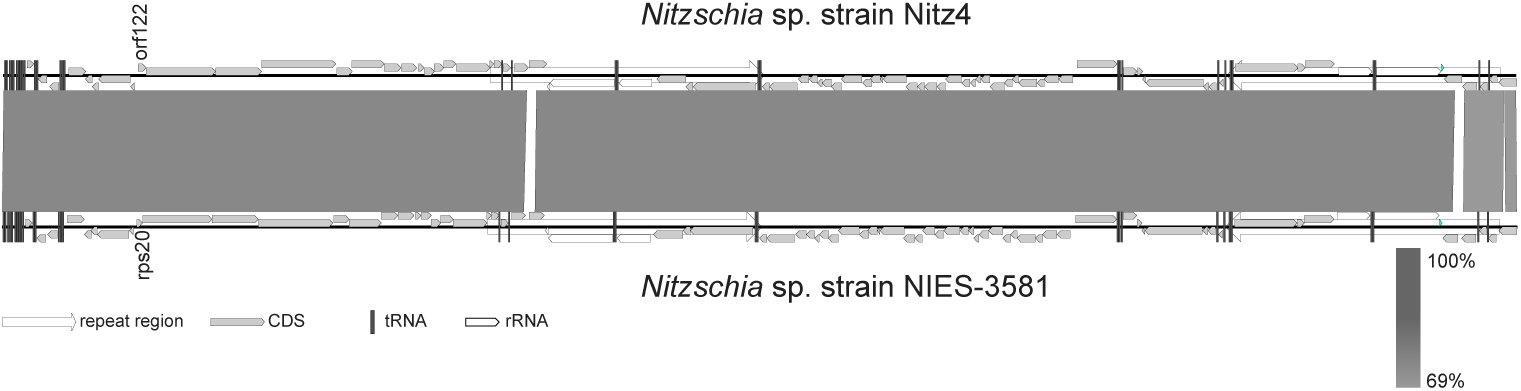
Conserved synteny, gene content, and sequence in the plastid genomes of two non-photosynthetic diatoms in the genus *Nitzschia*, Nitz4 (this study) and NIES-3581 (Kamikawa et al. 2015a). Unlabeled genes are shared between the two species.

## Discussion

### Monophyly of apochloritic diatoms

The small number of species (roughly 20) and narrow taxonomic range (all Bacillariales, mostly all *Nitzschia*) of apochloritic diatoms suggests that these species—the only known nonphotosynthetic diatoms—are monophyletic, reflecting a single loss of photosynthesis and transition to obligate heterotrophy in their common ancestor. Previous phylogenetic tests of this hypothesis were equivocal, however, with the plastid 16S rDNA sequences supporting monophyly of apochloritic species and the nuclear 28S d1–d2 gene splitting them, albeit with low bootstrap support, into three separate clades (Kamikawa et al. 2015b). Further underscoring the uncertainty in their relationships, the non-normalized, highly AT-rich plastid 16S sequences may have resulted in long-branch artifacts (Steel et al. 1993, Galtier and Gouy 1995), and the nuclear 28S dataset could not reject monophyly of apochloritic species (Kamikawa et al. 2015b), pointing to insufficient signal in this relatively short sequence fragment to address this question. As efforts to understand the causes and genomic consequences of loss of photosynthesis in diatoms accelerate (Kamikawa et al. 2015a, 2015b, 2017), it is important to determine if the shift occurred one or multiple times. The number, pattern, and timing of shifts has important implications for understanding the ease of transition(s) to obligate heterotrophy, the ecological setting of the transition(s), and, in practical terms, the power of comparative approaches to resolve these questions—as evolutionary replication, represented in this case by multiple losses of photosynthesis—maximizes the power of comparative methods to reveal possible mechanisms underlying these kinds of evolutionary transitions (e.g., Maddison and FitzJohn 2015). A single transition would, by contrast, greatly limit the power of comparative methods to uncover the underlying genomic and ecological drivers of the switch away from auto- or mixotrophy to obligate heterotrophy.

To address this problem, we increased both the number of taxa and genes available for phylogenetic analyses. We also normalized the plastid 16S rDNA sequences to guard against artefactual grouping of apochloritic species with highly AT-rich plastid genomes. Although support was generally low, the best trees inferred from genes representing all three genetic compartments uniformly supported monophyly of apochloritic *Nitzschia* (Fig. 2). In short, the best available data support a single loss of photosynthesis in *Nitzschia* and, by extension, diatoms as a whole. In light of this, future research efforts can focus attention away from questions that naturally arise for characters that evolve multiple times (e.g., the role of convergent evolution) and focus more on firmly placing the apochloritic clade within the broader Bacillariales phylogeny to help identify, for example, the ecological and mixotrophic characteristics of their closest relatives. Considerable divergence and phylogenetic structure exists within the apochloritic clade, with the largest combined-data tree (mito-lsu-new-ncbi) splitting taxa into two major subclades (Figs. 2 and S1). The modest level of species diversity, in turn, might make it feasible to sample each subclade more-or-less exhaustively and apply comparative approaches that will be useful for understanding the genomic and metabolic consequences of the switch to heterotrophy, including, for example, the rate of decay of the photosynthetic apparatus. All of this requires a robust, time-calibrated phylogeny of Bacillariales, which is one of the largest and most species-rich taxonomic orders of diatoms (Kociolek et al. 2018). This and other phylogenetic studies have begun resolving some parts of the Bacillariales tree using small numbers of commonly used phylogenetic markers (e.g., Lundholm et al. 2002, Rimet et al. 2011, Smida et al. 2014), but many relationships remain unresolved. With most of the traditional markers now more-or-less exhausted, effort should turn to much deeper sampling of the nuclear genome, which appears to hold great promise for resolving relationships across many phylogenetic scales within diatoms (Parks et al. 2018).

### The ecology and biogeography of apochloritic diatoms

Although relatively few apochloritic *Nitzschia* species have been formally described and named, the amount of phylogenetic diversity described in this (Fig. 2) and other studies (Kamikawa et al. 2015b), suggests that this clade could easily comprise on the order of 20 species. This seems even more probable when considering the relatively modest historical efforts to collect and characterize apochloritic diatoms. One emerging, and quite striking, theme among the small number of studies that focused on characterizing species diversity in this group is the large number and diversity of apochloritic *Nitzschia* that co-occur over very small spatial scales (e.g., within a sample from a single site). The diversity is apparent from both morphological and molecular phylogenetic evidence alike (Figs. 1 and 2; Blackburn et al. 2009a, Kamikawa et al. 2015b). Although some of the apochloritic taxa in our trees came from culture collections (Table 1 and Fig. S1), most of the taxa derive from a small number of mangrove-dominated habitats in Japan (Kamikawa et al. 2015b) and the United States (Figs. 1 and 2; Blackburn et al. 2009a). Both of these sites host multiple (apparently undescribed) apochloritic *Nitzschia* species that span the full phylogenetic breadth of the clade (Fig. S1). Focused efforts to collect and culture apochloritic *Nitzschia* from amenable habitats (e.g., seagrasses and mangroves) worldwide will show whether this anecdotal trend holds. If it is upheld, the next natural step will be to determine whether species co-occurrence is made possible by the sheer abundance of local resources in these habitats or rather by resource partitioning among species, either in fine-scale microhabitats or through specialization on different carbon sources. Of course, both of these alternatives require knowledge of the organic carbon species used by these species. Nevertheless, the phylogeny clearly shows species are not clustered geographically (Fig. S1), which suggests that many or most apochloritic diatom species have broad, presumably worldwide, geographic distributions and that the local species pool at any one site is the product of dispersal, not in situ speciation.

### Plastid genome reduction in apochloritic diatoms

Although probably different species, the two *Nitzschia* strains with sequenced plastid genomes belong to the same subclade (Fig. S1A, bottom) and so, in the context of the entire apochloritic clade, are very close relatives. Their close relationship was also evident in their plastid genomes, which were highly similar in nearly all respects. Additional sampling across the entire apochloritic clade will show whether all species share the same fundamental, highly reduced plastid genome—indicative of rapid genomic streamlining following the loss of photosynthesis in their common ancestor—or whether decay of the plastid genome is ongoing in some taxa. Similar to some green algae (Knauf and Hachtel 2002), diatoms have retained a nearly full set of ATP synthase genes whose products, instead of functioning in ATP hydrolysis, generate a proton gradient for tat-dependent protein translocation across the thylakoid membrane (Kamikawa et al. 2015a). This model underscores both the compartmentalized nature of carbon metabolism in diatoms (Smith et al. 2012, Kamikawa et al. 2017) and, consequently, the indispensable nonphotosynthetic plastids in diatoms.

In addition to understanding the specific consequences for the plastid genome to following the loss of photosynthesis, a fuller understanding of these plastid genomes can shed light on photosynthetic diatom plastids as well. For example, the retention several conserved ORFs in these nonphotosynthetic genomes—in the context of near wholesale loss of the photosynthetic apparatus—strongly suggests these conserved, diatom-specific ORFs have functions that are unrelated, or only peripherally related, to photosynthesis. These ORFs include ycf41, ycf89, and ycf90. Finally, the loss of photosynthesis can have cascading effects on rates of sequence evolution all three genomes (Nickrent et al. 1998). Branch lengths based on plastid, mitochondrial, and nuclear genes showed that the AT-driven rate acceleration in the plastid genomes of apochloritic species appears to be restricted to the plastid genome alone (Fig. 2). This does not rule out that there have been other effects on the mitochondrial and nuclear genomes including, for example, gene content and whether some of the missing plastid genes have been transferred to the nucleus.

### Conclusions

The discovery that apochloritic *Nitzschia* are monophyletic represents an important step forward in understanding the loss of photosynthesis and switch to obligate heterotrophy in diatoms—a transition that has occurred just once in a lineage of some 100,000 species of photoautotrophs. A clearer view of this relationship highlights new research avenues and priorities, including more intensive taxon sampling within Bacillariales to identify the closest relatives of the apochloritic clade, which almost certainly evolved from a mixotrophic ancestor (Hellebust and Lewin 1977, Figueroa-Martinez et al. 2015). The exact carbon sources, modes of carbon uptake and utilization, and the degree of carbon specialization within and across species remains unclear for photosynthetic and nonphotosynethetic *Nitzschia* alike. Bacillariales features a diverse set of taxa with fascinating biology, motivating the development of excellent genomic resources for this group (Basu et al. 2017, Kamikawa et al. 2017, Mock et al. 2017). A nuclear genome sequence for a nonphotosynthetic *Nitzschia* could help address several of these outstanding questions, leading to hypotheses that can be directly tested in a group of diatoms that has proven highly amenable to experimental manipulation (e.g., Lewin and Lewin 1967, Azam and Volcani 1974, McGinnis and Sommerfeld 2000).

## Acknowledgments

We thank Sarah Hamsher, Nina Lundholm, and Pat Kociolek for early discussions on the phylogeny of Bacillariales. This work was supported by National Science Foundation (NSF) (grant number DEB-1353131) and multiple awards from the Arkansas Biosciences Institute to AJA. This research used computational resources available through the Arkansas High Performance Computing Center, which were funded through multiple NSF grants and the Arkansas Economic Development Commission.

## Supporting Information

**Table S1.**
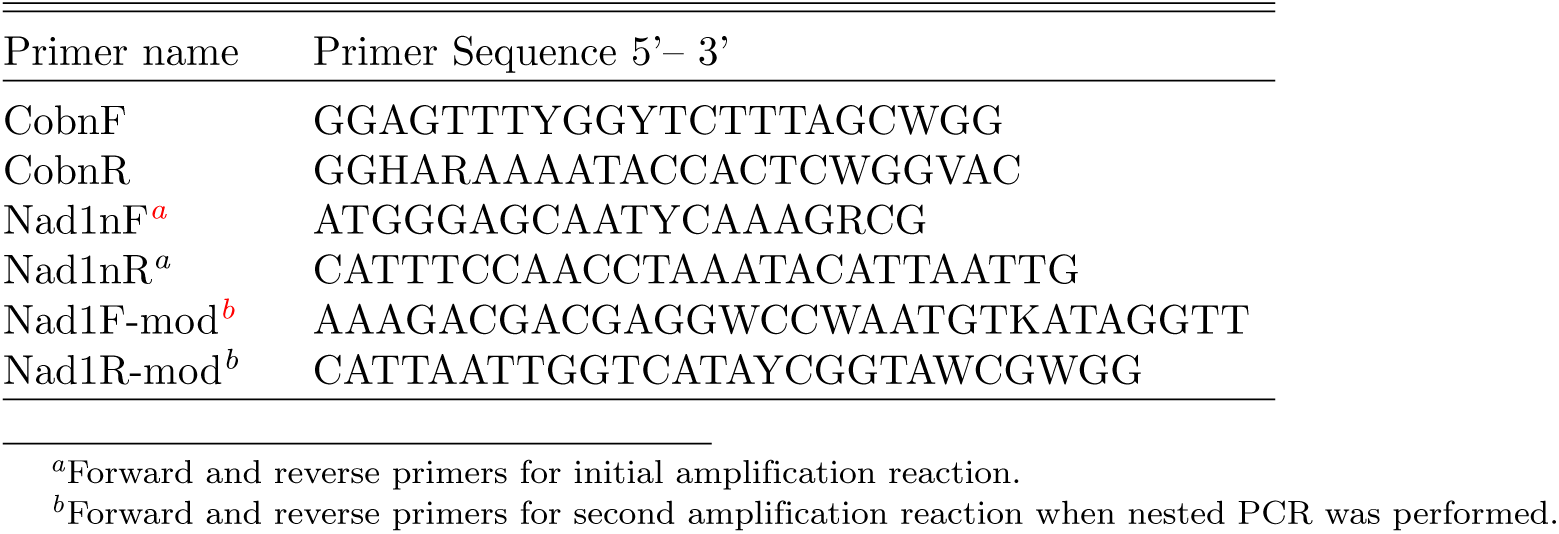
Primers used to amplify and sequence *cob* and *nad1* fragments for study taxa.

**Table S2.** Overlapping and adjacent genes in the plastid genomes of *Nitzschia* sp. nitz4 and *Nitzschia* sp. NIES-3581. Groups of overlapping or adjacent genes are separated by empty rows with the coordinates showing the degree of overlap. All but two groups are shared between the two genomes.

| *Nitzschia* sp. nitz4 |  |  |  | *Nitzschia* sp. NIES.3581 |  |  |  |
| --- | --- | --- | --- | --- | --- | --- | --- |
| Gene | Coordinates | over<sup>a</sup> | adj<sup>b</sup> | Gene | Coordinates | over | adj |
| rpl19 | complement(3679..3993) |  |  | rpl19 | complement(3681..3995) |  |  |
| orf96 | complement(3986..4276) | + |  | urf97 | complement(3970..4260) | + |  |
| atpB | complement(4267..5694) | + |  |  |  |  |  |
| orf122 | 6041..6409 |  |  |  |  |  |  |
| rpoB | 6406..9513 | + |  |  |  |  |  |
| rpoC1 | 9517..11580 |  |  | rpoC1 | 9410..11479 |  |  |
| rpoC2 | 11577..14900 | + |  | rpoC2 | 11476..14820 | + |  |
| rps2 | 14938..15627 |  |  | rps2 | 14858..15547 |  |  |
| sufB | 15624..17072 | + |  | sufB | 15544..16992 | + |  |
| sufC | 17072..17815 |  | + | sufC | 16992..17735 |  | + |
| atpG | 18859..19311 |  |  | atpG | 18777..19229 |  |  |
| atpF | 19298..19732 | + |  | atpF | 19216..19650 | + |  |
| atpA | 20300..21805 |  |  | atpA | 20206..21720 |  |  |
| rpl35 | 21783..21977 | + |  | rpl35 | 21714..21908 | + |  |
| rps12 | complement(36097..36471) |  |  | rps12 | complement(36024..36398) |  |  |
| rpl31 | complement(36471..36680) |  | + | rpl31 | complement(36398..36604) |  | + |
| rps9 | complement(36687..37091) |  |  | rps9 | complement(36612..37016) |  |  |
| rpl13 | complement(37088..37537) | + |  | rpl13 | complement(37013..37462) | + |  |
| rpoA | complement(37534..38466) | + |  | rpoA | complement(37459..38391) | + |  |
| rps11 | complement(38487..38891) |  |  | rps11 | complement(38412..38822) |  |  |
| rps13 | complement(38830..39273) | + |  | rps13 | complement(38755..39198) | + |  |
| secY | complement(39409..40455) |  |  | secY | complement(39334..40398) |  |  |
| rps5 | complement(40427..40930) | + |  | rps5 | complement(40370..40873) | + |  |
| rps17 | complement(43317..43556) |  |  | rps17 | complement(43288..43527) |  |  |
| rpl29 | complement(43546..43782) | + |  | rpl29 | complement(43517..43756) | + |  |
|  |  |  |  | dnaK | 48083..49930 |  |  |
|  |  |  |  | tRNA-Arg | 49930..50003 |  | + |
| TOTAL |  | 12 | 2 |  |  | 10 | 3 |

<sup>a</sup>Overlapping
<sup>b</sup>Adjacent

**Figure S1.**
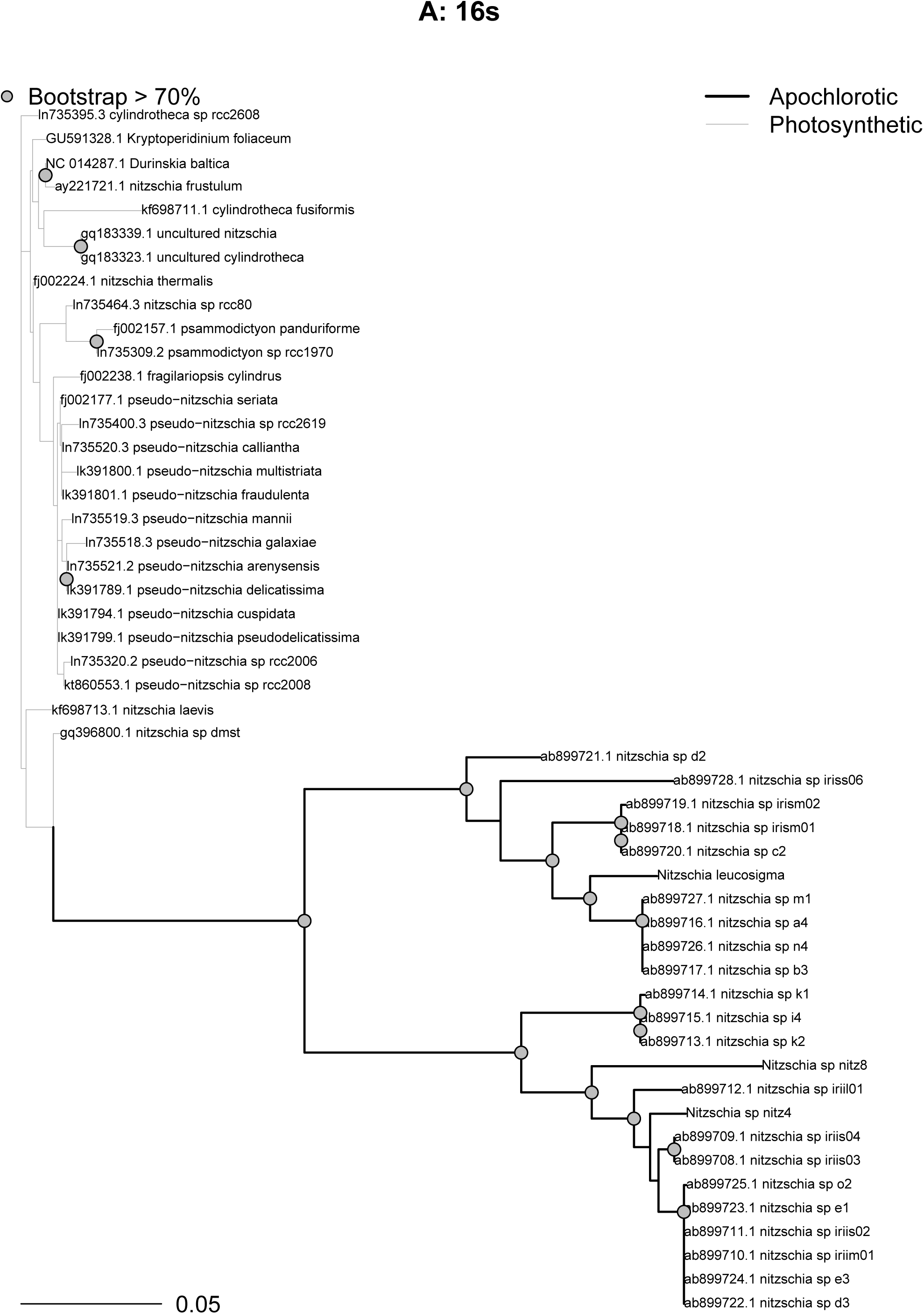
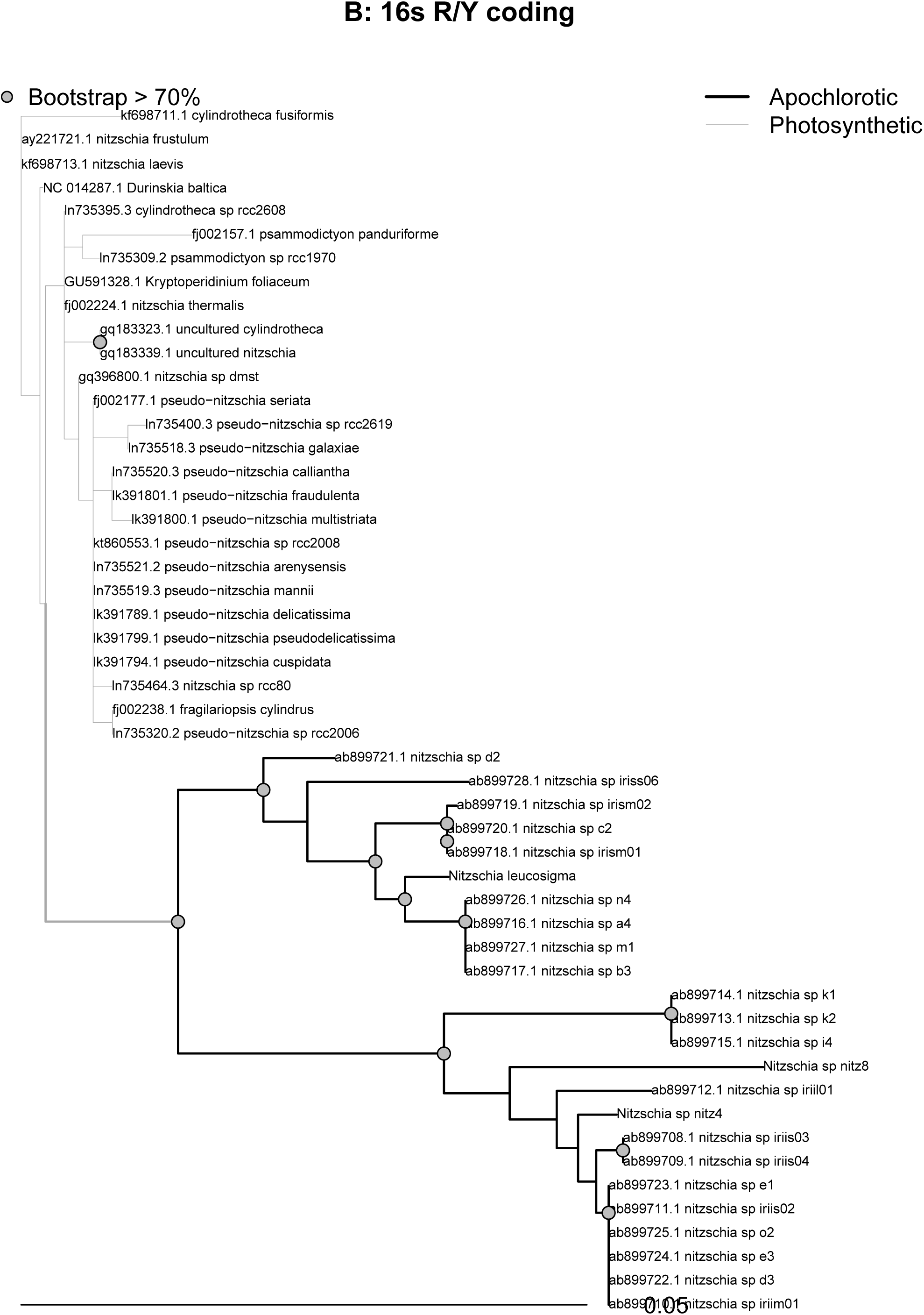
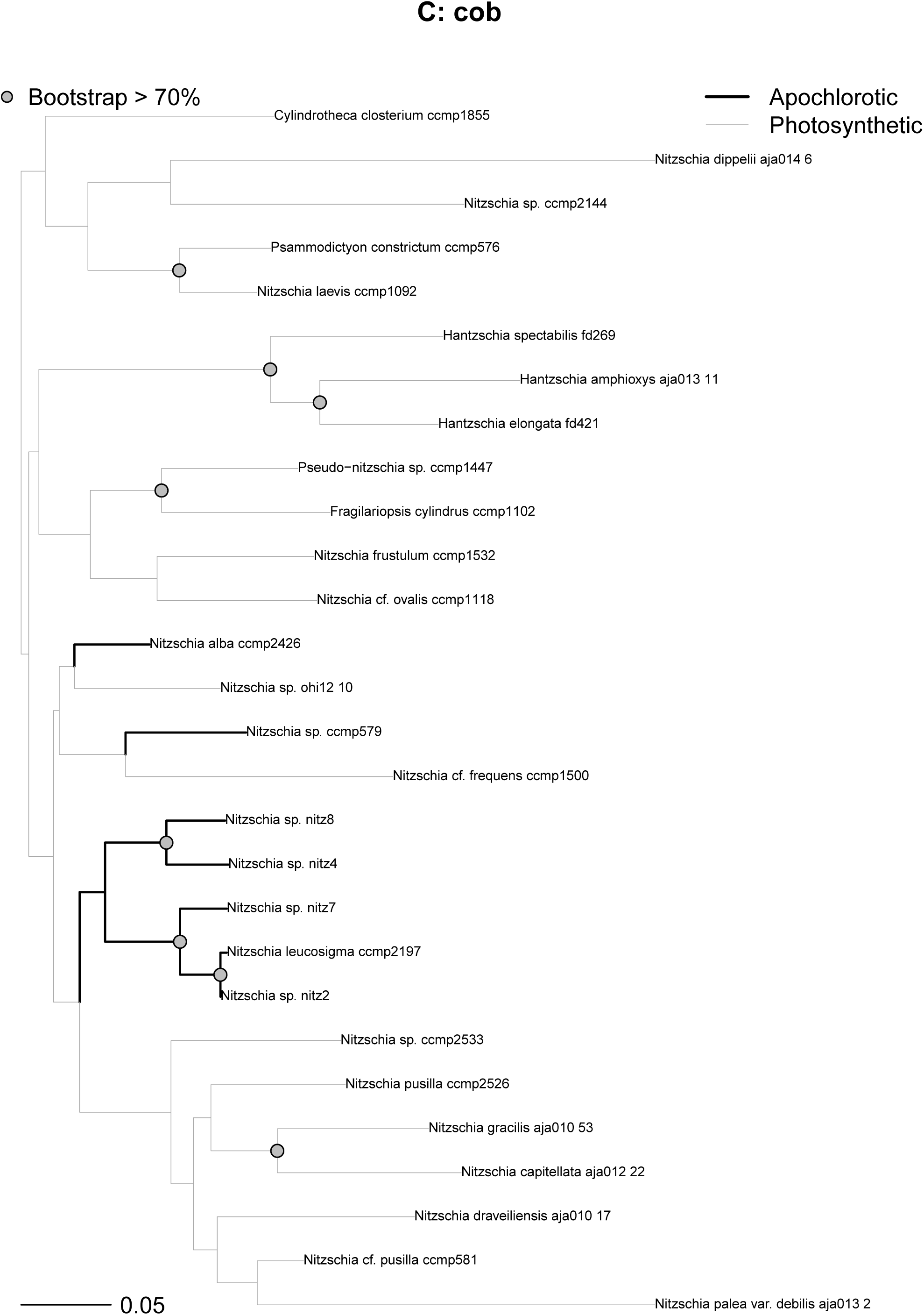
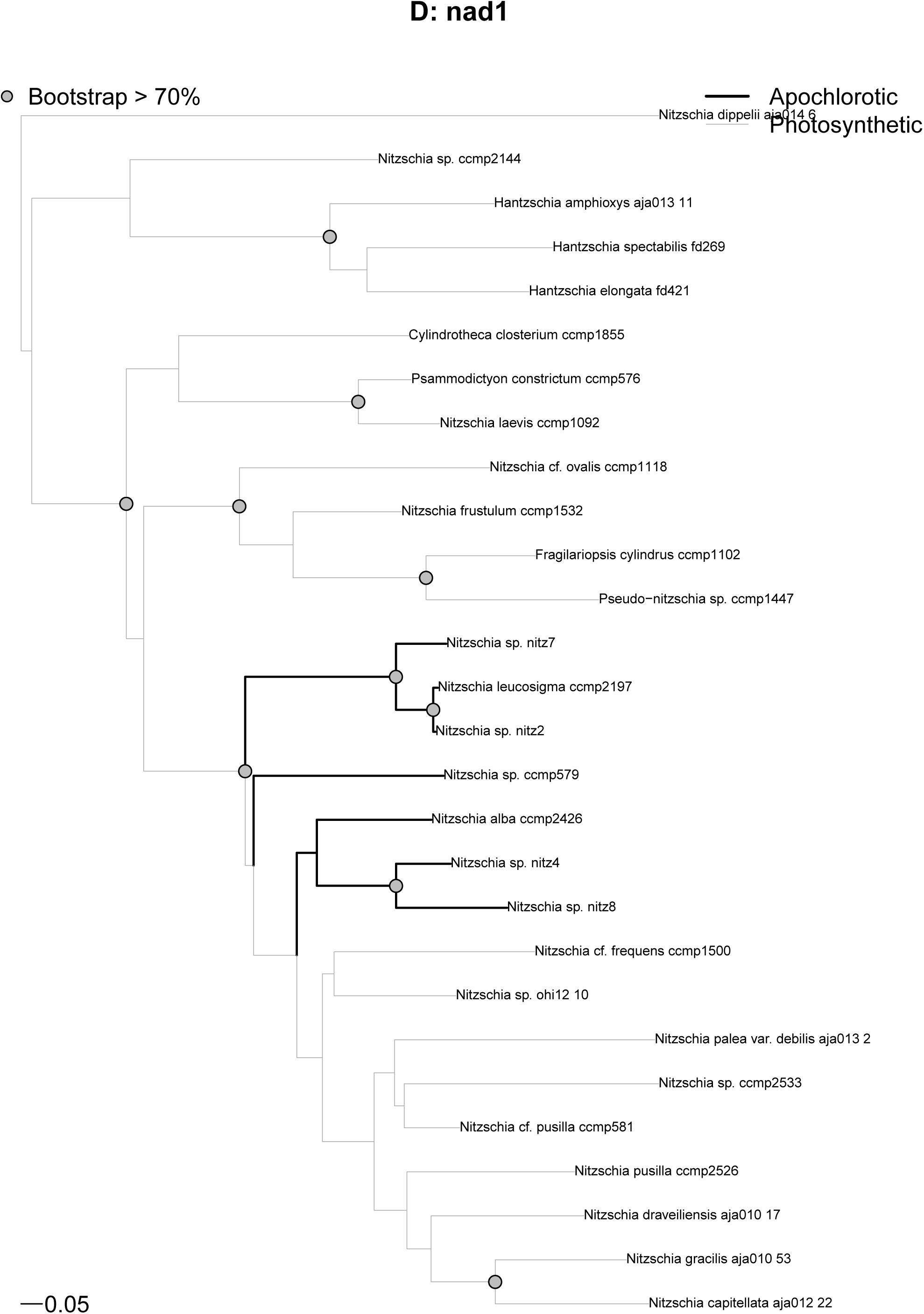
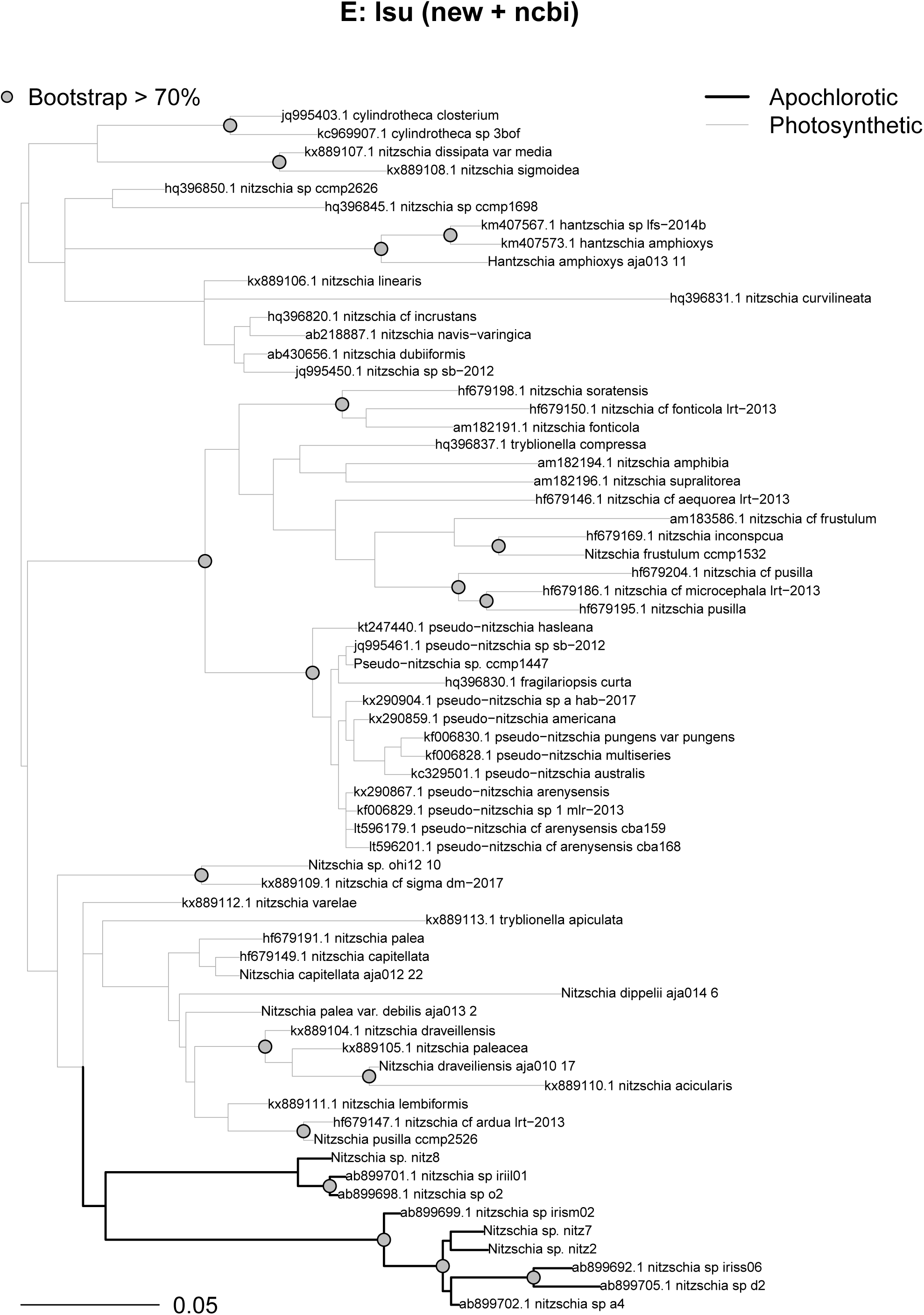
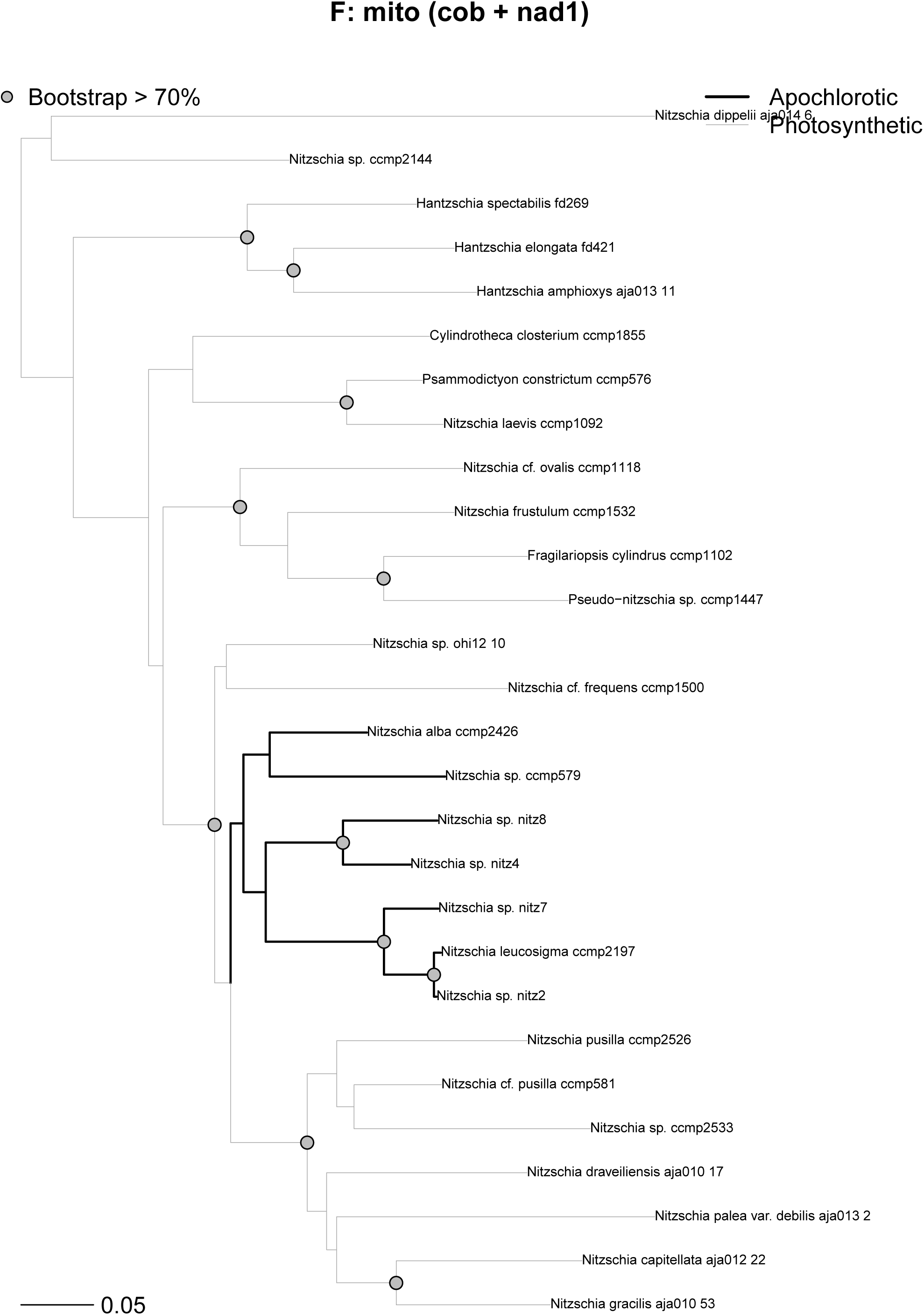
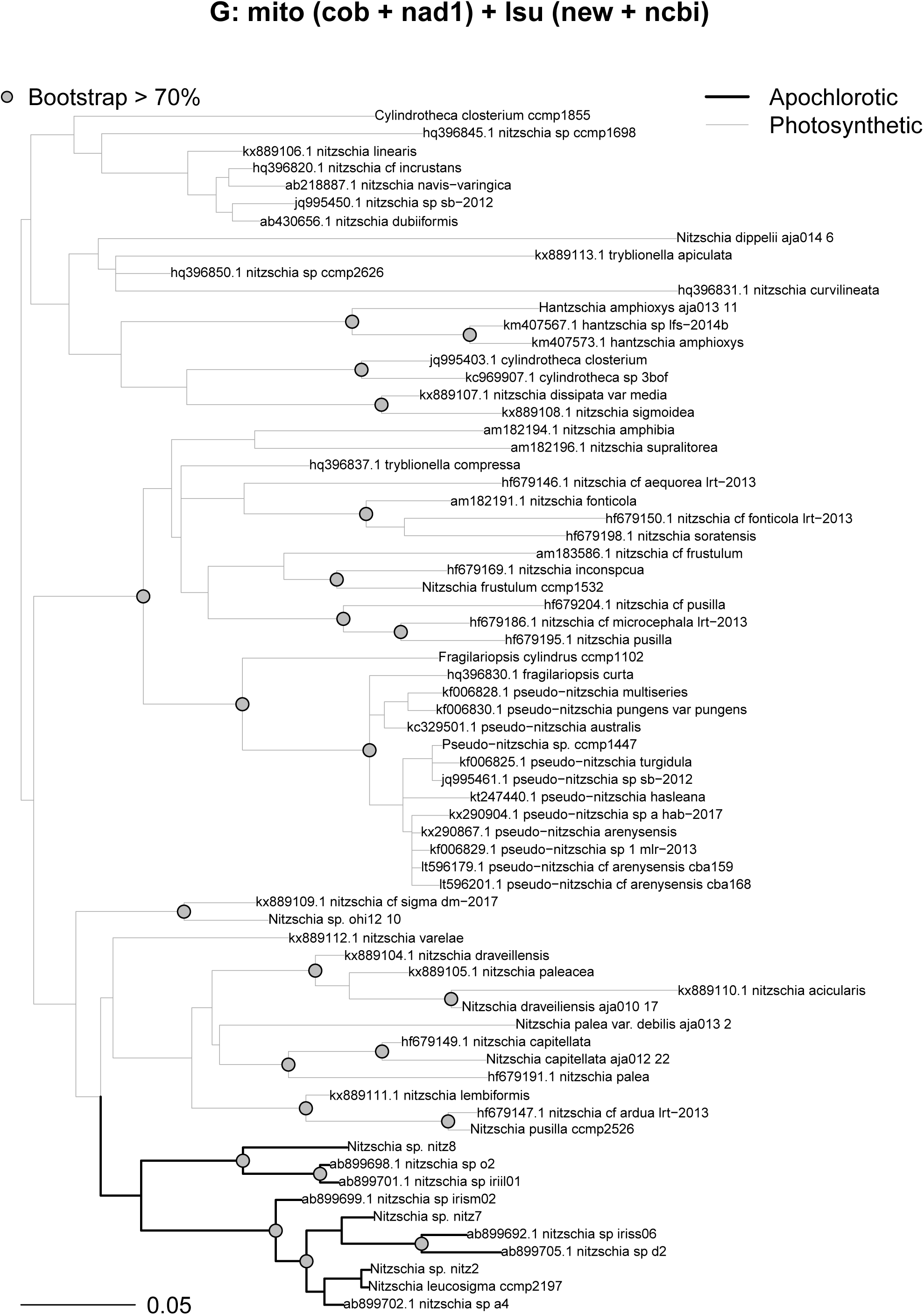
Phylogenetic trees for plastid 16S (A), plastid 16S transformed into purine/pyrimidine (R/Y) coding (B), mitochondrial *cob* (C), *nad1* (D), a densely sampled 28S d1–d2 matrix with sequences from this study and GenBank (E), *cob* and *nad1* combined (F), and a large combined nuclear and mitochondrial gene matrix with sequences from this study and GenBank (G). In the 16S tree, *Nitzschia* sp. NIES-3581 is identified by its synonym, iriis04.

